# Agronomic, physiological and molecular characterization of rice mutants revealed key role of ROS and catalase in high temperature stress tolerance

**DOI:** 10.1101/739433

**Authors:** Syed Adeel Zafar, Amjad Hameed, Muhammad Ashraf, Abdus Salam Khan, Zia-ul-Qamar, Xueyong Li, Kadambot H.M. Siddique

## Abstract

Plants adapt to harsh environments particularly high temperature stress by regulating their physiological and biochemical processes, which are key tolerance mechanisms. Thus, identification of heat-tolerant rice genotypes and reliable selection indices are crucial for rice improvement programs. Here, we evaluated the response of a rice mutant population for high-temperature stress at the seedling and reproductive stages based on agronomic, physiological and molecular traits. The estimate of variance components revealed significant differences (*P*<0.001) among genotypes, treatments and their interaction for almost all traits. Principal component analysis showed significant diversity among the genotypes and traits under high-temperature stress. The mutant ‘HTT-121’ was identified as the most heat tolerant mutant with higher grain yield, panicle fertility, cell membrane thermo-stability (CMTS) and antioxidant enzyme levels under heat stress conditions. Various seedling-based morpho-physiological traits (leaf fresh weight, relative water contents, malondialdehyde, CMTS) and biochemical traits (superoxide dismutase, catalase and hydrogen peroxide) explained variations in grain yield that could be used as selection indices for heat tolerance in rice at early growth stages. Notably, heat sensitive mutants showed a significant accumulation of ROS level, reduced activities of catalase and upregulation of *OsSRFP1* expression under heat stress, suggesting their key role in regulating heat tolerance in rice. The heat-tolerant mutants identified in this study could be used in breeding programs and the development of mapping populations to unravel the underlying genetic architecture for heat-stress adaptability.

**Summary text for table of contents:** Heat stress probably due to changing climate scenario has become a serious threat for global rice production. On the other side, efforts to develop high yielding cultivars have led to the reduced genetic variability to withstand harsh environmental conditions. This study aimed to identify novel heat tolerant mutants developed through gamma irradiation which will provide a unique genetic resource for breeding programs. Further, we have identified reliable selection indices for screening heat-tolerant rice germplasm at early growth stages.

## Introduction

Climate change has emerged as a major challenge worldwide, affecting human health, agricultural production and natural resources, among others (Piao *et al.* 2010). One of the major effects of climate change is the onset of high-temperature stress, which will threaten global food security (Hasegawa *et al.* 2018). To address these issues, modern breeding programs have reoriented their aims to focus on stress factors (Borém *et al.* 2012). To attain genetic gain, breeding programs need genetic variants from which to choose, select and introgress adaptation attributes, i.e., heat tolerance or other parameters to assist in dealing with climate fluctuations. In relation to abiotic stresses, breeding has developed cultivars suitable for areas where the crops were not adapted previously (Tester and Langridge 2010).

Rice is a major staple food crop that sustains the lives of about three billion people around the world (Krishnan *et al.* 2011). Rice production needs to increase by 50% by 2030 to fulfill the global population of rice-dependent countries (Ahmadi *et al.* 2014). Climate change has already influenced many aspects of rice production, including yield reduction (Garrett *et al.* 2014). Rice productivity in the 21st century will encounter unprecedented challenges due to changing climatic conditions, including unstable patterns of precipitation and temperature. Rice production is very susceptible to high-temperature stress as it results in poor seed set due to pollen sterility or anther indehiscence (Arshad *et al.* 2017). High temperature increases membrane injury and impairs metabolic functions which affect agronomic traits directly linked to yield (Mohammed and Tarpley 2009; Zafar *et al.* 2018). The mechanisms of heat-stress tolerance in plants are complex and governed by many genes, proteins, antioxidants and other factors that involve various physiological and biochemical amendments in cells, such as modifications to cell membrane function and structure and primary and secondary metabolites (Huang and Xu 2008). At the onset of stress, the plasma membrane is one of the first components affected, and its stability under stressed conditions is regarded as the main indicator of heat tolerance in crop plants (Blum and Ebercon 1981). Similarly, chlorophyll content is used to evaluate the physiological status of crop plants under abiotic stress (Lichtenthaler *et al.* 2000). High temperature also induces the over-accumulation of reactive oxygen species (ROS) that can cause cell injury via programmed cell death (Xu *et al.* 2006). To overcome the damaging effects of higher ROS levels, plants produce antioxidants as a tolerance mechanism (Kumar *et al.* 2012). Combining different stress-tolerance parameters at different developmental stages assists in the development of cultivars tolerant to a multitude of stress factors (Fleury *et al.* 2010). In this context, seedling resistance can be instrumental for later stage development as well as being important at the specified stage (Ayalew *et al.* 2015; Rehman *et al.* 2016). However, studies relating seedling-stage resistance with reproductive-stage heat tolerance are scanty, particularly in rice.

In the past few decades, the focus on developing high-yielding rice cultivars has narrowed the genetic diversity of rice, particularly for traits related to biotic and abiotic stresses. To address this key issue, we aimed to develop a mutant rice population that could provide beneficial alleles as a resource for breeding. Mutant germplasm resources have been developed for other crops that offer better parental combinations and speed up breeding programs. The present study included 39 mutants of cv. Super Basmati, and IR-64 as a heat-sensitive check, under normal and heat-stress conditions to identify mutants with heat tolerance at both the seedling and reproductive stages to pinpoint useful and reliable heat-tolerance indicators. The identified heat-tolerant mutants will serve as a useful genetic resource for further genetic studies and breeding for heat-tolerant rice.

## Materials and methods

The germplasm comprised 41 rice genotypes including cv. Super Basmati (approved basmati rice variety in Pakistan), 39 mutants (M5 generation) of Super Basmati developed by gamma irradiation (using doses of 20–30 Grey), and rice cultivar ‘IR-64’ as a heat-sensitive check (Poli *et al.* 2013) (Supplemental Table S1). The mutants were developed at the Nuclear Institute for Agriculture and Biology (NIAB), Faisalabad, Pakistan.

### Screening for physiological and biochemical traits at the seedling stage

Two sets of seeds were sown in plastic pots filled with an equal quantity of clean soil under controlled conditions in a growth chamber at normal temperature (28±2°C). Both sets were sown in triplicate (15 seedlings per replicate) and placed in the dark until seedling emergence (3–4 days). After emergence, a 12 h photoperiod (irradiance of 120 μmol m^−2^ s^−1^) was maintained. After 14 days, one set of uniform seedlings was subjected to heat stress (45±2°C) for 12 h in a growth chamber running at 45±2°C under the same light conditions mentioned above, while the other set remained at normal temperature and served as the control. After high-temperature exposure, the seedlings had a three-day recovery period at normal temperature; after which, leaf samples (first and second leaf) from five seedlings were collected from each replicate for physiological and biochemical analysis and immediately stored at –80°C until further use.

### Trait measurements

The fresh weight of leaves and seedlings were measured immediately after harvest to avoid water loss. The turgid weight of leaves was recorded after soaking in water for 24 h. The dry weight of leaves and seedlings were recorded after drying in a 90°C oven for 36 h.

### Cell membrane thermo-stability (CMTS)

CMTS was calculated using the method of Martineau *et al.* (1979):

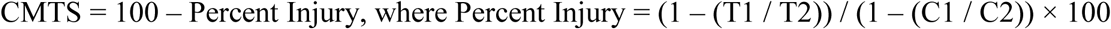

where T1 and T2 refer to the first and second conductivity measurement (after autoclaving), respectively, of heat-stressed leaf segments and C1 and C2 refer to the first and second conductivity measurement, respectively, of control plant leaf segments.

### Relative water content (RWC)

RWC was calculated using the formula of Yamasaki and Dillenburg (1999):

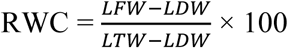

where LFW, LDW and LTW are leaf fresh, dry and turgid weight, respectively. SFW and SDW in the text hereafter refer to fresh and dry weight of seedlings, respectively.

### Malondialdehyde (MDA)

The level of lipid peroxidation in leaf tissue was measured in terms of MDA content using the method of Heath and Packer (1968) with minor modifications as described by Dhindsa *et al.* (1981).

### Chlorophyll contents

Chlorophyll *a* and *b* concentrations were determined following the method of Arnon (1949) and carotenoid concentration determined following the method of Davies (1976). Absorbance of the extract was measured at 663, 645, 505, 470 and 453 nm using a spectrophotometer (HITACHI, U2800).

To estimate biochemical parameters including total soluble proteins (TSP), enzymatic and non-enzymatic antioxidants and other stress biomarkers, leaves (0.15 g) were homogenized in 1.5 ml potassium phosphate buffer (pH 7.4) using a cold mortar and pestle. Samples were centrifuged at 14,462 *g* for 10 min at 4°C. The supernatant was separated and used to determine enzyme activities and other biochemical assays as described below.

### Superoxide dismutase (SOD)

SOD activity was assayed using the method of Giannopolitis and Ries (1977). The reaction solution (1 ml) comprised double distilled water (400 µl), 200 mM potassium phosphate buffer pH 7.8 (250 µl), 13 mM methionine (100 µl), Triton X (100 µl), NBT (50 µl), 1.3 µM riboflavin (50 µl), and 50 µl enzyme extract. The test tubes containing the reaction solution were irradiated (15 W fluorescent lamps) at 78 µmol m^−2^ s^−1^ for 15 min. The absorbance of the irradiated solution was determined at 560 nm. One unit of SOD activity was defined as the amount of enzyme that caused 50% inhibition of photochemical reduction of NBT.

### Catalase (CAT)

CAT activity was estimated using the method of Sizer (1952). The assay solution (3 ml) contained 50 mM phosphate buffer (pH 7.0), 59 mM H_2_O_2_, and 0.1 ml enzyme extract. The decrease in absorbance of the reaction solution at 240 nm was recorded every 20 s. An absorbance change of 0.01 min^−1^ was defined as 1 U of CAT activity. Enzyme activities were expressed on a fresh weight basis.

### Peroxidase (POD)

POD activity was measured using the method of Chance and Maehly (1955) with some modifications. The assay solution (1 ml) contained distilled water (545 µl), 50 mM phosphate buffer (250 µl) (pH 7.0), 20 mM guaiacol (100 µl), 40 mM H_2_O_2_ (100 µl), and 5 µl enzyme extract. The reaction was initiated by adding the enzyme extract. The increase in absorbance of the reaction solution at 470 nm was recorded every 20 s. One unit of POD activity was defined as an absorbance change of 0.01 min^−1^.

### Ascorbate Peroxidase (APX)

APX activity was measured using the method of Dixit *et al.* (2001). The assay buffer was prepared by mixing 200 mM potassium phosphate buffer (pH 7.0), 10 mM ascorbic acid, and 0.5 M EDTA. The assay solution contained assay buffer (1 ml), H_2_O_2_ (1 ml), and 50 µl supernatant. The oxidation rate of ascorbic acid was estimated by following the decrease in absorbance at 290 nm every 30 s (Chen and Asada 1989).

### Total phenolics content (TPC)

A microcolorimetric method, as described by Ainsworth and Gillespie (2007), was used Folin– Ciocalteu (F–C) reagent for the total phenolics assay. A standard curve was prepared using different concentrations of gallic acid, and a linear regression equation was calculated. Phenolic content (gallic acid equivalents) of samples was determined using the linear regression equation.

### Protease

For protease estimation, leaves were homogenized in a medium comprising 50 mM potassium phosphate buffer (pH 7.8). Protease activity was determined by casein digestion assay as described by Drapeau (1974). Using this method, 1 U is the amount of enzyme that releases acid-soluble fragments equivalent to 0.001 A280 per minute at 37°C and pH 7.8. Enzyme activity was expressed on a fresh weight basis.

### Esterases

The α-esterases and β-esterases were determined according to the method of Van Asperen (1962) using α-naphthyl acetate and β-naphthyl acetate as substrates, respectively. The reaction mixture consisted of substrate solution [30 mM α or β-naphthyl acetate, 1% acetone, and 0.04 M phosphate buffer (pH 7)] and enzyme extract. The mixture was incubated for exactly 15 min at 27°C in the dark, then 1 ml of staining solution (1% fast blue BB and 5% SDS mixed in a ratio of 2:5) was added followed by incubation for 20 min at 27°C in the dark. The amount of α- and β-naphthol produced was measured by recording the absorbance at 590 nm.

### Total soluble protein (TSP)

Estimation of quantitative protein was executed using the method of Bradford (1976) by mixing 5 µl of supernatant and 95 µl NaCl (150 mM) with 1.0 ml of dye reagent [0.02 g Coomassie Brilliant Blue G-250 dye dissolved in 10 ml 95% ethanol and 20 ml 85% (w/v) phosphoric acid, and diluted to 200 ml]. The mixture was allowed to sit for 5 min to form a protein-dye complex before recording the absorbance at 595 nm.

### Total oxidant status (TOS)

TOS was determined using a novel method formulated by Erel (2005) based on the oxidation of the ferrous ion to the ferric ion. The assay mixture contained reagent R1 (stock xylenol orange solution (0.38 g in 500 μL of 25 mM H_2_SO_4_), 0.49 g NaCl, 500 μL glycerol made up to 50 mL with 25 mM H_2_SO_4_), sample extract, and reagent R2 (0.0317 g θ-anisidine, 0.0196 g ferrous ammonium sulfate II). After 5 min, absorption was measured at 560 nm.

### Hydrogen peroxide (H_2_O_2_)

ROS was measured in terms of H_2_O_2_ following the instructions provided by hydrogen peroxide assay kit (Beyotime, China). Briefly, 100 mg leaf tissue was extracted with 1 ml 50 mM sodium phosphate buffer (pH 7.4) and centrifuged for 15 min at 12000 *g* at 4 °C. The supernatant was used to measure OD at 560 nm. H_2_O_2_ was then estimated from standard curve.

### RNA isolation and quantitative real time PCR

RNA was extracted from contrasting heat tolerant and sensitive mutants along with cv. Super basmati and IR-64 at three time points; before heat stress (designated as control), 24 h after heat stress (designated as 24-HAS), and after three days of recovery (called RC). RNAPrep Pure Plant kit (TIANGEN, China) was used to isolate total RNA. 1 µg RNA was reverse transcribed into cDNA using HiScript II Q RT Supermix (Vazyme). ChamQ SYBR qPCR Master Mix was used for the qPCR reaction using an ABI Prism 7500 sequence detection system with the programs recommended by the manufacturer. *ACTIN1* gene was used as an internal control. qRT-PCR primer sequences for *SODA*, *SODB*, *CATA*, *CATB, OsSRFP1* and *Actin1* are listed in Table S2.

### Field evaluations

The same set of genotypes were evaluated under natural field conditions in 2014 under two temperature scenarios (normal and high-temperature stress) for various yield-contributing agronomic traits including plant height (PH), number of productive tillers per plant (PTP), panicle length (PL), number of spikelets per main panicle (SMP), panicle fertility percentage (PF), thousand-grain weight (TGW) and grain yield per plant (PY). For normal temperature conditions, the material was grown in the field at NIAB, Faisalabad, Pakistan (in northern Punjab province, 31.41° N, 73.07° E). For HTS, the same material was grown in district Multan, Pakistan (in southern Punjab province, 30.19° N, 71.46° E) which is usually warmer than Faisalabad. Uniform fields were prepared at both locations to minimize environmental variation for soil properties. Water and fertilizer were applied according to the recommendations of the agriculture department of Punjab, Pakistan. Each field was divided into three plots, with each plot treated as one replicate. All 39 mutants, along with the heat-sensitive check and cv. Super Basmati, were grown in each plot using plant-to-plant and row-to-row distances of 20 cm to avoid any shading effect on neighboring plants (Poli *et al.* 2013). Each mutant was sown in five rows of six plants in each replicate. At physiological maturity, six to eight representative plants from the middle rows of each replicate were selected for agronomic data measurements to avoid confounding border effects (Chaturvedi *et al.* 2017). The data for each recorded parameter were average across replicates.

Based on the overall performance of the mutants in the agronomic and physiological evaluation conducted in 2014, selected heat-tolerant and heat-sensitive mutants along with parent cv. Super Basmati and sensitive check IR-64 were evaluated in 2016 under controlled HTS conditions to validate their heat tolerance. For the control treatment, plants were grown under natural field conditions, and for the HTS treatment, plants at the start of anthesis were covered with a polythene sheet during the day (3 m above ground, serving as a tunnel) for ten days to impose heat stress. A difference of 4–6°C between treatments was recorded during the heat stress. Two sides of the tunnel (facing each other) were left open for air flow to maintain the same humidity level as the outside (∼70%). The same agronomic traits were measured as for 2014.

### Heat susceptibility index

The heat susceptibility index for grain yield (HSI-GY) was calculated using the formula [(1 – Y / Yp) / D] as described by Khanna-Chopra and Viswanathan (1999) where Y and Yp are the yield of a genotype under heat stress and normal conditions, respectively, and D (stress intensity) = (1 – X / Xp) where X and Xp are the mean of Y and Yp, respectively.

### Statistical analysis

Data were analyzed using analysis of variance (ANOVA) to test the significance of genotypes, environments and their interaction (G × E) on the studied plant traits using SAS version 9.2 (SAS Institute, Cary, NC, USA). Principal component analysis (PCA) was performed using XL-STAT software (version 2014), and Pearson’s correlation was performed using the corrplot package in R software. Agronomic data from 2014 (control and HTS treatment) were used for all statistical analysis unless otherwise stated. Mean data with standard errors for 2014 and 2016 are presented in Figures 4 and 5.

## Results

### Temperature scenario of field trials and crop growth

The daily mean and maximum temperature during the 2014 growing period was obtained from the Pakistan meteorological department and is presented as Supplemental Figure S1. Multan (designated HTS environment) recorded an overall increasing trend in mean and maximum temperatures, relative to Faisalabad (designated normal environment). At both locations, crops started active anthesis and pollination from 15 August, with seed set in mid-September. Anthesis and fertilization are the most critical and sensitive stages of rice growth for temperature stress. During this time in 2014 (18-20 August), differences of 2.7–4.7°C were observed between the normal and HTS environments. In 2016, differences of 4–6°C were observed during anthesis under control and HTS. The maximum temperature during anthesis in the HTS treatment in the tunnel ranged from 38.4 to 42.7°C while the maximum temperature outside ranged from 34.3 to 37.6°C in 2016. Thus, a relatively high temperature was observed in the heat-stress treatment in the field environment in Multan in 2014 as well as the tunnel in 2016.

### Genotype performance under normal and HTS conditions

The ANOVA displayed highly significant differences (*P*<0.001) for genotypes, environments and G × E for most traits (Table 1). There were a few non-significant relationships, including the effect of environment on carotenoids and TGW, and the effect of G × E on PL and TGW, so these traits would not be useful selection indicators for heat tolerance in rice.

**Table 1.** Mean square values from the analysis of variance for the effect of genotype, environment and their interaction on various morpho-physiological, biochemical and agronomic traits. Significance level: *P < 0.05, **P < 0.01, ***P < 0.001.

| SOV | Replications | Genotypes (G) | Environments (E) | G × E | Error |
| --- | --- | --- | --- | --- | --- |
| df | 2 | 40 | 1 | 40 | 162 |
| LFW | 0.76 | 17.55*** | 434.03*** | 7.23*** | 0.91 |
| LDW | 0.04 | 0.77*** | 15.39*** | 0.39*** | 0.02 |
| RWC | 45.74 | 196.62** | 8905.90*** | 179.12* | 59.84 |
| SFW | 35.7 | 129.81*** | 2568.67*** | 33.49*** | 1.16 |
| SDW | 0.12 | 7.40*** | 99.30*** | 6.73*** | 0.46 |
| MDA | 6.4 | 295.86*** | 2680.41*** | 247.48*** | 2.89 |
| Lyco | 1.58 | 14.33*** | 14.18*** | 11.68*** | 0.43 |
| chl *a* | 30.06 | 3104.19*** | 1100.18*** | 2395.00*** | 11.38 |
| chl *b* | 14.16 | 48894.98*** | 386.66*** | 34263.53*** | 7.13 |
| Car | 1.59 | 40.28*** | 0.09 | 45.54*** | 0.82 |
| TCC | 62.11 | 38456.45*** | 811.12** | 29627.52*** | 65.25 |
| TSP | 2.11 | 1602.44*** | 69001.76*** | 1404.90*** | 6.38 |
| CAT | 65.13 | 50414.40*** | 155823.92*** | 43251.41*** | 148.19 |
| POD | 69516.67 | 576552967.41*** | 42627828673.47*** | 452875124.83*** | 368793.33 |
| APX | 487.5 | 168642.65*** | 1250.00** | 156330.00*** | 143.89 |
| SOD | 18.11 | 938.62*** | 4332.74*** | 1394.45*** | 20.91 |
| Prot | 5254.17 | 381716.42*** | 34237812.50*** | 282310.62*** | 4655 |
| Estr | 2573.07* | 137365.05*** | 15964012.28*** | 102739.93*** | 630.42 |
| TPC | 73266.67 | 16284778.08*** | 37380363.52*** | 23116462.68*** | 64728.89 |
| TOS | 10.74 | 115.52*** | 3069.81*** | 209.29*** | 3.77 |
| PH | 1.23 | 578.98*** | 5099.78*** | 75.23*** | 5.91 |
| PTP | 0.14 | 20.31*** | 129.15*** | 10.18*** | 0.13 |
| PL | 0.52 | 7.88** | 10.84* | 2.73 | 1.81 |
| SMP | 0.22 | 673.22*** | 551.26*** | 438.18*** | 7.08 |
| PF | 2.42 | 47.32*** | 419.67*** | 24.29*** | 2.36 |
| TGW | 0.05 | 7.10*** | 0.96 | 0.17 | 0.24 |
| PY | 2376.85 | 851669.44*** | 426122.74*** | 231776.10*** | 2229.61 |

**Abbreviations:** df, degrees of freedom; LFW, leaf fresh weight; LDW, leaf dry weight; RWC, relative water content; SFW, seedling fresh weight; SDW, seedling dry weight; MDA, malondialdehyde; Lyco, lycopene; chl *a*, chlorophyll *a*; chl *b*, chlorophyll *b*; Car, carotenoids; TCC, total chlorophyll content; TSP, total soluble proteins; CAT, catalase; POD, peroxidase; APX, ascorbate peroxidase; SOD, superoxide dismutase; TPC, total phenolic content; TOS, total oxidant status; PH, plant height; PTP, productive tillers per plant; PL, panicle length; SMP, spikelets per main panicle; PF, panicle fertility; TGW, thousand grain weight; PY, paddy yield (grain yield).

### Principal component analysis revealed genetic diversity among mutants

A genotype-trait (G-T) biplot was developed using PCA to observe genetic diversity among the evaluated genotypes for traits under both normal and HTS environments (Figure 1). In the normal environment, the first 10 PCs had eigenvalues >1 and contributed 78.45% of the cumulative variability (Table S3). A G-T biplot was constructed using the first two PCs (PC1 and PC2), which accounted for 15.96% and 12.77% of individual variability, respectively (Figure 1). Similarly, in the HTS environment, the first 10 PCs had eigenvalues >1 and contributed 81.34% of the cumulative variability (Table S3), with PC1 and PC2 accounting for 14.24% and 12.65% of individual variability, respectively (Figure 1).

**Fig 1.**
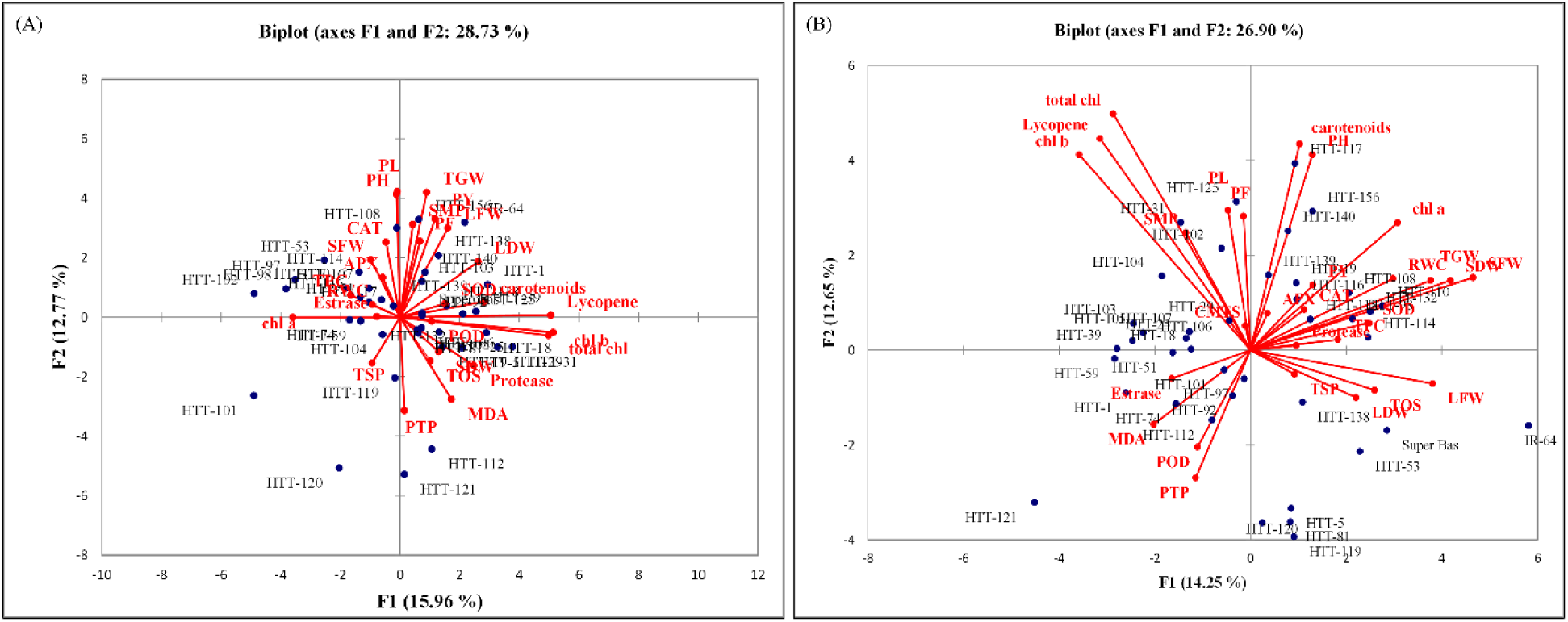
Principal component analysis showing biplot for genotypes and studied traits under normal (A) and high-temperature stress (B).

In the normal environment, PC1 was mainly represented by chlorophyll *b*, lycopene, total chlorophyll content, carotenoids, LDW, protease, MDA, LFW, SOD, SDW, PY, POD, TOS, TGW and PF, while PC2 was mostly characterized by PL, TGW, PH, PY, SMP, LFW, PF, CAT, SFW, LDW, APX and TPC (Table S4). In the HTS environment, PC1 was mainly represented by SFW, SDW, LFW, TGW, chlorophyll *a*, RWC, TOS, SOD, LDW, TPC, PY, PH, CAT, carotenoids, protease and TSP, while PC2 was primarily characterized by total chlorophyll content, lycopene, carotenoids, PH, chlorophyll *b*, PL, PF, chlorophyll *a*, SMP, SFW, RWC, SDW, TGW, PY, CAT and APX (Table S4).

The biplot analysis indicated that under normal conditions, mutants HTT-120, HTT-121, HTT-112 and HTT-101, and traits PTP, MDA, PL, PH and TGW were largely dispersed and away from the origin and had high genetic variability (Figure 1). Similarly, in the HTS environment, mutants HTT-121, IR64, HTT-119, HTT-81, HTT-120, HTT-5 and HTT-117 were highly dispersed and far away from the origin, which indicated high genetic variability and importance of these genotypes for selection. Mutant HTT-117 was very close to traits PH and carotenoids and showed higher phenotypic values for these traits. LFW, SFW, SDW, TGW, RWC, PH, chlorophyll a and carotenoids fall on the positive X-axis and were far away from the origin, which showed high variability and importance of these traits in the HTS environment. In addition, the biplot analysis showed that the studied genotypes and traits had higher genetic variability in the HTS environment than the normal environment.

### Correlation test revealed association among various traits

Pearson’s correlation analysis was performed using seedling-stage data of physiological and biochemical traits and reproductive-stage data of agronomic traits from 2014 to identify significant correlations among seedling-based and reproductive-stage-based traits with grain yield. The correlation analysis revealed significant positive and negative correlations among the studied traits, especially with grain yield (Figure 2). The analysis also showed an association of various seedling-based traits with different yield-related traits under both environments (normal and HTS). In the normal environment, LDW (*r =* 0.35), PH (*r =* 0.46), PL (*r =* 0.35) and TGW (*r =* 0.49) had significant (*P*<0.05) positive correlations with yield (PY), and LFW (*r =* 0.54), LDW (*r =* 0.32), SFW (*r =* 0.42), PH (*r =* 0.35) and PL (*r =* 0.49) had significant (*P*<0.05) positive correlations with TGW. In the HTS environment, protease (*r =* 0.36) and PF (*r =* 0.32) had significant positive correlations with PY, and protease (*r =* 0.32) and PL (*r =* 0.45) had significant positive correlations with TGW. The number of productive tillers per plant (PTP) is an important agronomic trait for breeding. in the normal environment, PTP had significant negative correlations with PF (*r =* – 0.48), SMP (*r =* –0.44) and CAT (*r =* –0.46). In the HTS environment, PTP also had significant negative correlations with these traits (PF, SMP, and CAT) along with PH and RWC. In both environments, MDA (an indicator of oxidative damage) had significant negative correlations with CAT, POD and SOD.

**Fig 2.**
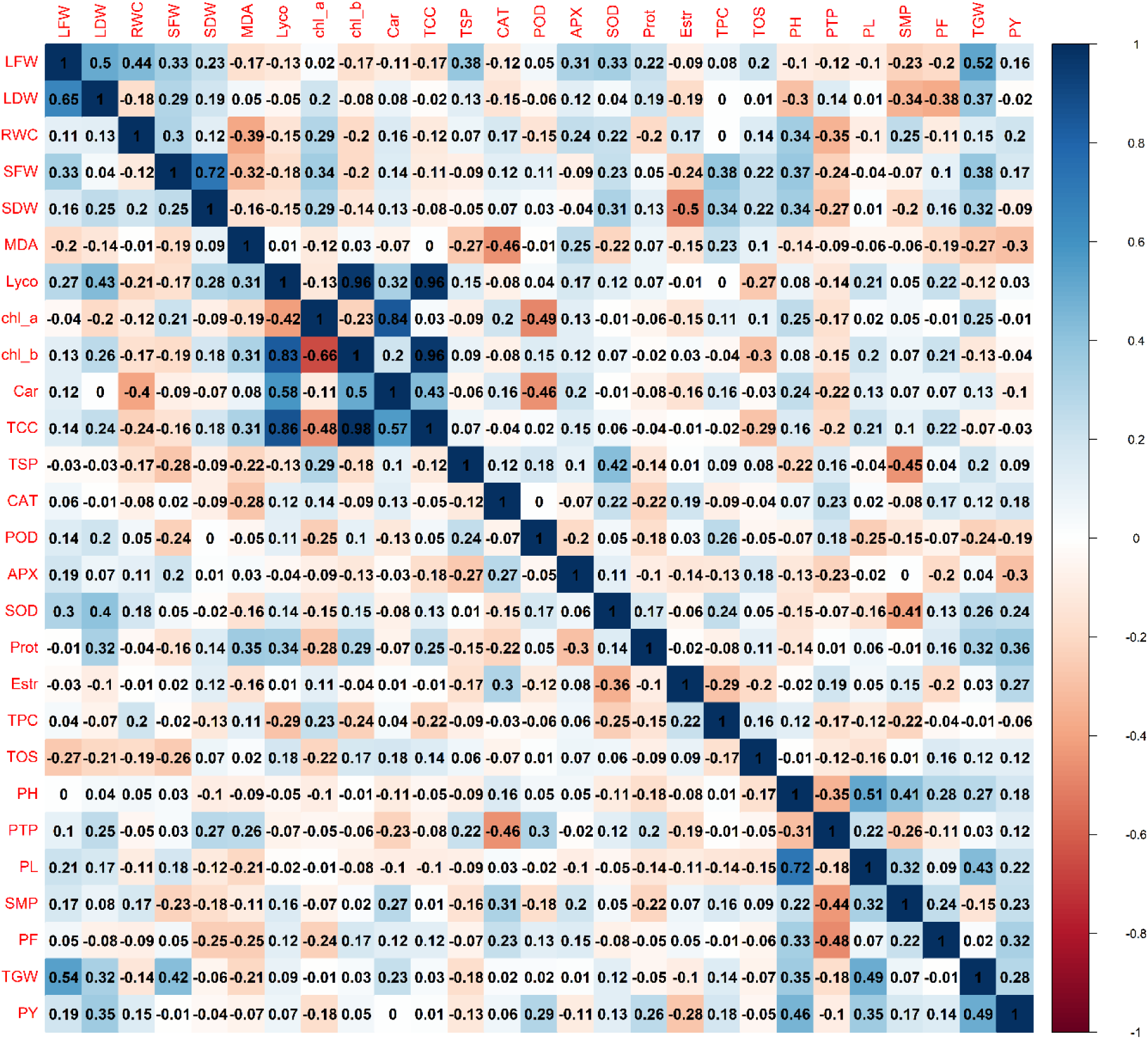
Correlation matrix showing Pearson’s correlation among traits in the rice mutant population under control (lower left diagonal) and high temperature (upper right diagonal). The scale bar on the right indicates the intensity of the correlation from 1 (highest positive in dark blue) to –1 (highest negative in red). **Abbreviations:** LFW, leaf fresh weight; LDW, leaf dry weight; RWC, relative water content; SFW, seedling fresh weight; SDW, seedling dry weight; MDA, malondialdehyde; Lyco, lycopene; chl *a*, chlorophyll *a*; chl *b*, chlorophyll *b*; Car, carotenoids; TCC, total chlorophyll content; TSP, total soluble proteins; CAT, catalase; POD, peroxidase; APX, ascorbate peroxidase; SOD, superoxide dismutase; Prot, protease; Estr, esterase; TPC, total phenolic content; TOS, total oxidant status; PH, plant height; PTP, productive tillers per plant; PL, panicle length; SMP, spikelets per main panicle; PF, panicle fertility; TGW, thousand-grain weight; PY, paddy yield (grain yield).

### Effect of HTS on grain yield

HSI-GY indicated the percent reduction in grain yield under HTS. Based on HSI-GY, the genotypes were divided into three groups *viz*. heat tolerant, moderately heat-tolerant and heat sensitive (Figure 3). Ten mutants were heat sensitive (HSI-GY > 7, mutants with >10% reduction in GY under HTS), 11 genotypes (including nine mutants) were moderately heat-sensitive (HSI-GY > 7, genotypes with <10% reduction in GY under HTS), and 20 mutants were heat tolerant (HSI-GY < 0, genotypes with no decline in GY under HTS). The heat-sensitive check (IR-64) and the parent of evaluated mutants (Super Basmati) were moderately heat-sensitive genotypes with almost 1% and 0.5% reductions in yield, respectively. Twenty-one genotypes (including Super Basmati) performed better than IR-64 in terms of grain yield.

**Fig 3.**
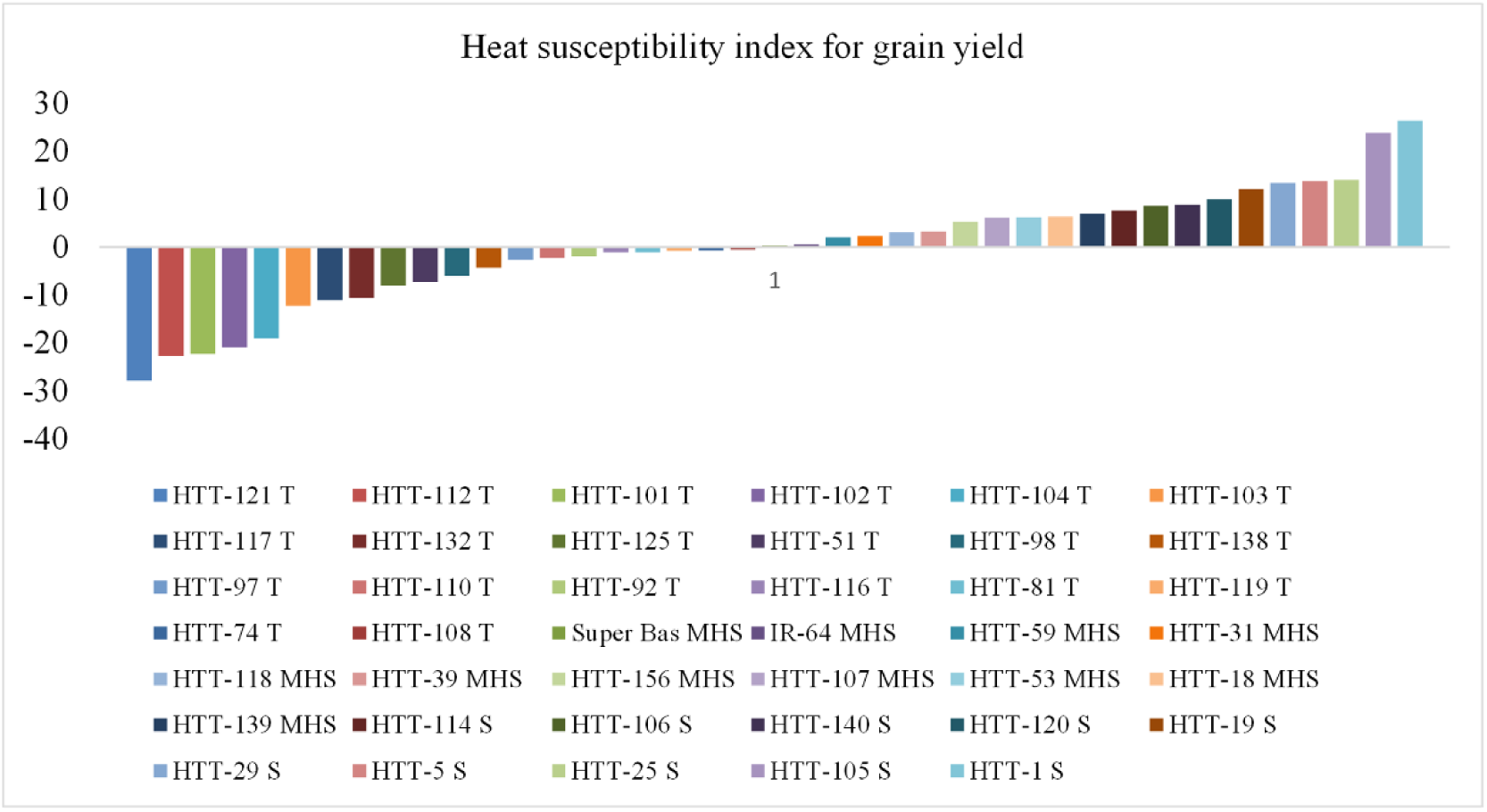
Heat susceptibility index for grain yield showing the degree of susceptibility to high temperature. T, MHS and S refers to tolerant, moderately tolerant and sensitive to high temperature.

### Mean performance of contrasting mutants for some agronomic, physiological and biochemical traits

Based on the HSI-GY from 2014, we evaluated the most heat-tolerant and least heat-tolerant (sensitive) mutants, along with the sensitive check (IR-64) and cv. Super Basmati, in 2016 under controlled temperature conditions to confirm the reproducibility and sustainability of data. The mean data for grain yield and panicle fertility from both years is presented in Figure 4A and 4B. In addition, data for those seedling-based morpho-physiological and biochemical traits that had significant associations with yield and TGW (LFW, RWC, SOD, CAT, MDA and CMTS) are presented in Figures 4C, 4D and 5A–D. A phenotypic comparison of panicles from HTT-121 (most heat-tolerant mutant), HTT-1 (least heat-tolerant mutant) and IR-64 (heat-sensitive check) under normal and HTS environments in 2016 is shown in Figure 6.

**Fig 4.**
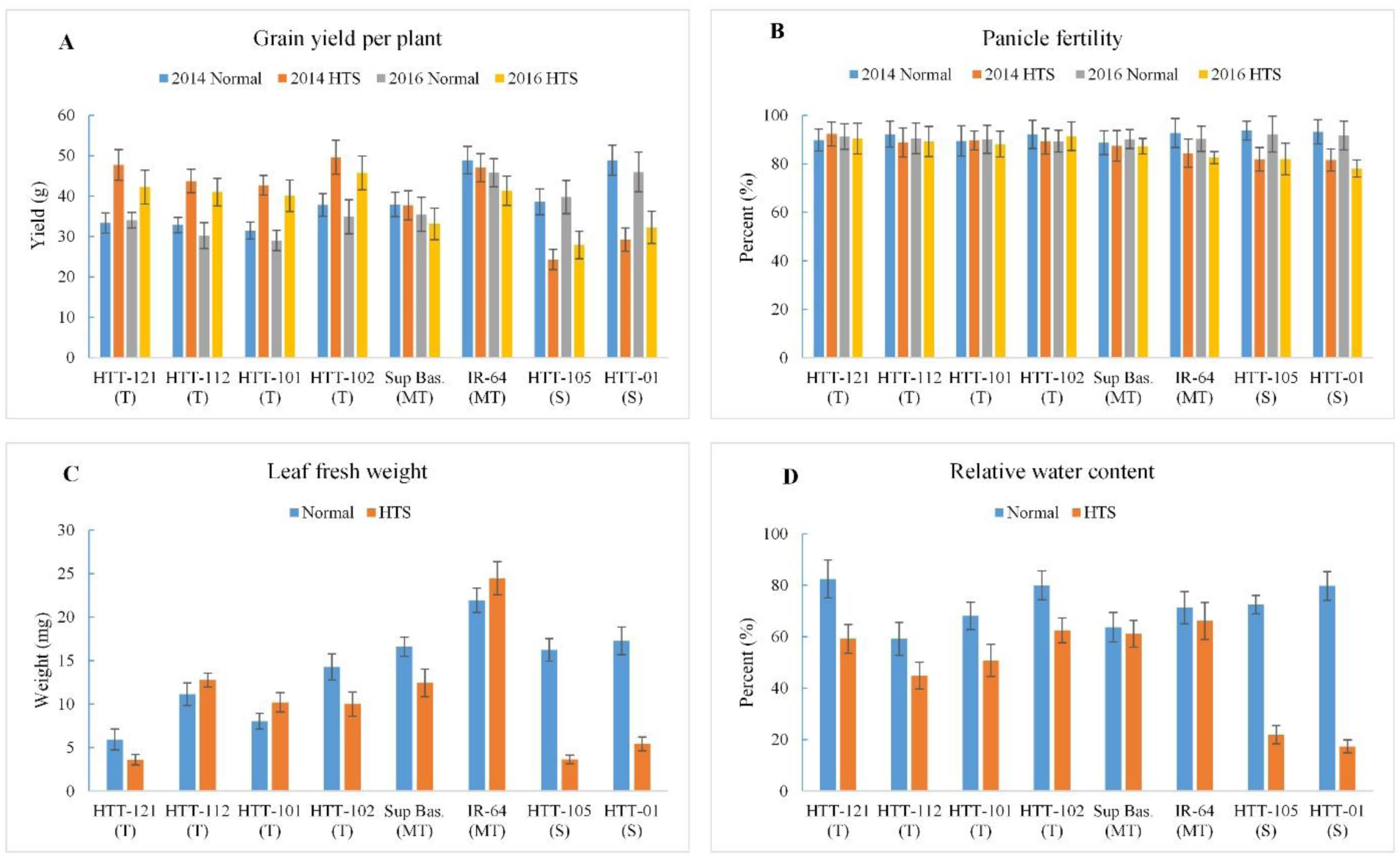
Effect of high temperature on grain yield (A), panicle fertility (B), leaf fresh weight (C) and relative water content (D). Values represent means ± SD.

**Fig 5.**
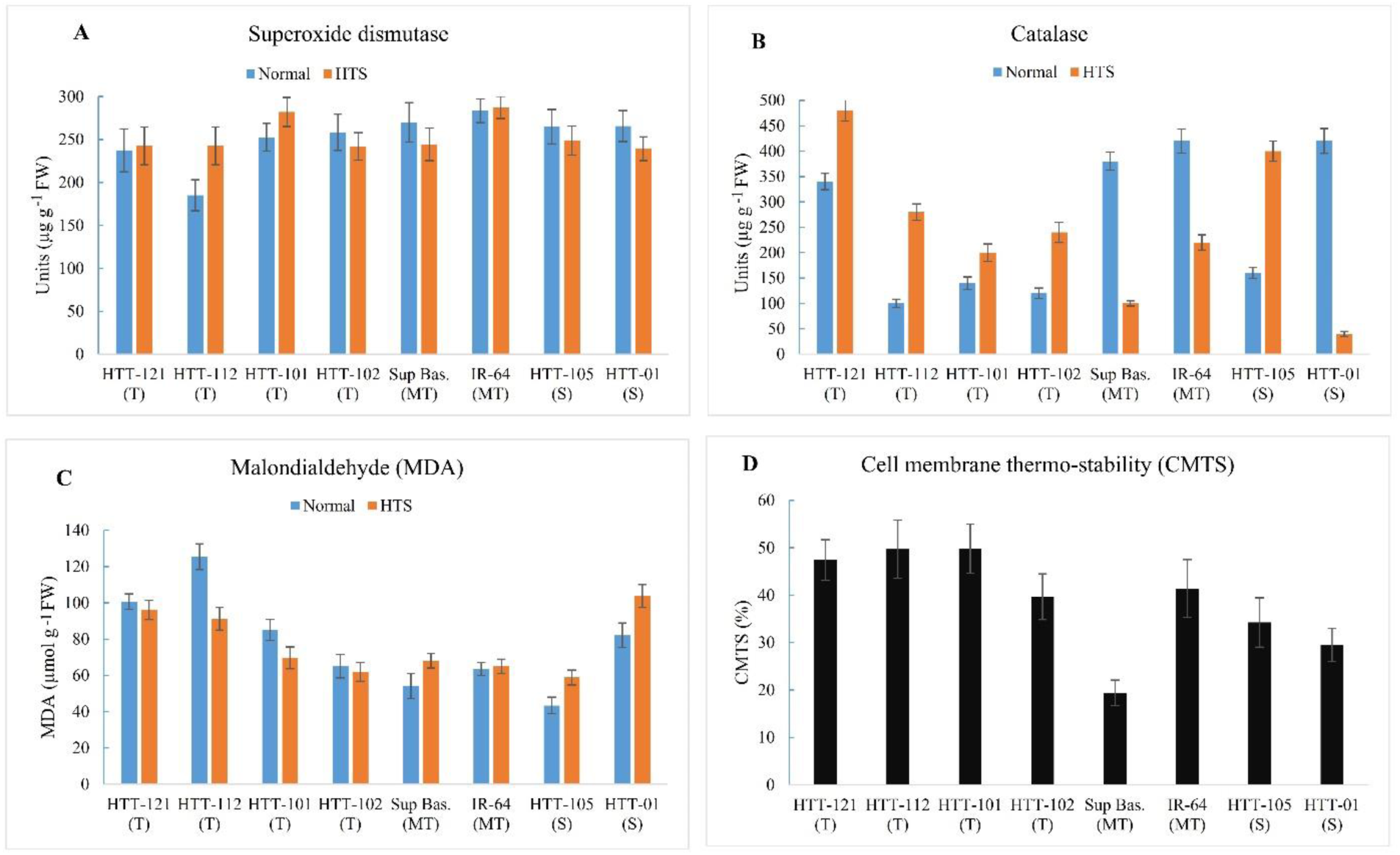
Effect of high temperature on the activity of superoxide dismutase (A), catalase (B), malondialdehyde (C) and cell membrane thermo-stability (D). Values represent means ± SD.

**Fig 6.**
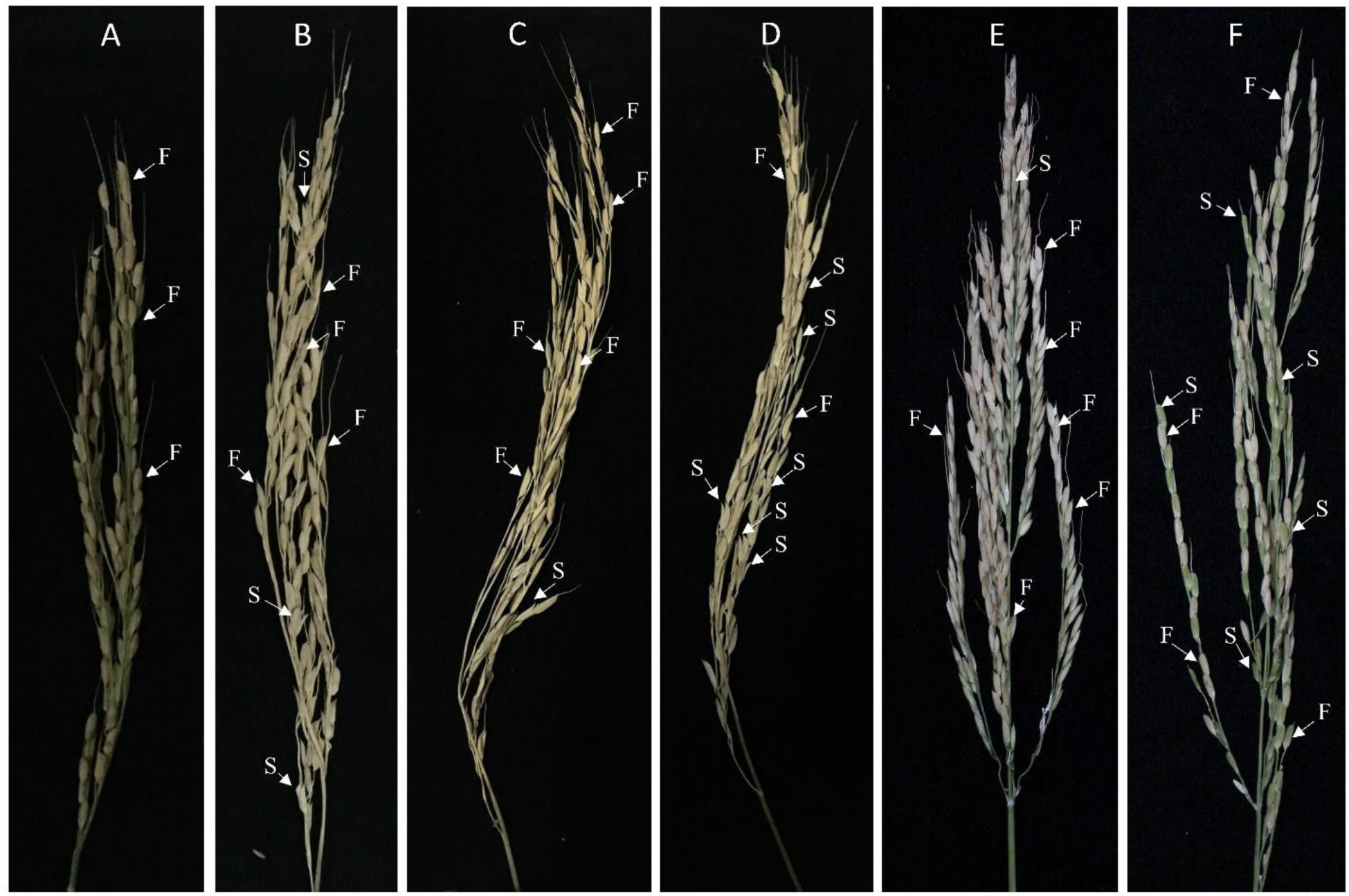
Comparison of panicle fertility under control and high-temperature stress. A, C and E represent panicles of HTT-121, HTT-1 and IR-64, respectively, under normal (control) condition. B, D and F represent panicles of HTT-121, HTT-1 and IR-64, respectively, under high-temperature stress. The F and S stand for fertile and sterile spikelets, respectively. Spikelets with open tips or green color represent sterile spikelets with no seed set.

The selected heat-tolerant mutants (HTT-121, HTT-112, HTT-101 and HTT-102) produced higher grain yields under HTS than the normal environment in both years (Figure 4A). The sensitive check (IR-64) and cv. Super Basmati had slightly lower yields under HTS than the normal environment. However, HTS significantly reduced grain yield in heat-sensitive mutants (HTT-1 and HTT-105) in both years. Similarly, HTS significantly reduced PF in heat-sensitive mutants (HTT-1 and HTT-105) in both years (Figure 4B). However, no significant differences in PF were observed in HTT-121, HTT-112, HTT-101 or HTT-102 under normal and HTS environments. The HTS significantly reduced LFW in HTT-1 and HTT-105, but any differences in the other mutants and IR-64 were not significant (Figure 4C). Similarly, RWC declined significantly in HTT-1 and HTT-105 under HTS (Figure 4D). Unexpectedly, HTS also significantly decreased RWC in HTT-121—ranked as tolerant among the mutants with better performance overall—but not as significantly as the heat-sensitive mutants.

Antioxidants such as SOD and CAT protect plant cells from the oxidative damage caused by abiotic stress by detoxifying ROS. The heat-tolerant mutants, apart from HTT-102, had higher SOD activity under HTS than the normal environment but the reverse was the case for the heat-sensitive mutants (Figure 5A). Similarly, HTS induced CAT activity in HTT-121, HTT-112, HTT-101 and HTT-102 (heat-tolerant mutants) but significantly reduced SOD activity in HTT-1 (most heat-sensitive mutant) and IR-64 (Figure 5B). However, HTT-105 maintained higher CAT activity under HTS, which may be a compensatory response of its defense system. MDA, which shows membrane lipid peroxidation, is an indicator of oxidative damage caused by higher ROS levels. Lower MDA levels were observed in all heat-tolerant mutants (HTT-121, HTT-112, HTT-101 and HTT-102) under HTS than the normal environment (Figure 5C). In contrast, HTS increased MDA levels in the heat-sensitive mutants (HTT-105 and HTT-1) and Super Basmati. CMTS estimates the level of cell injury in mutants (Figure 5D). Overall, the heat-tolerant mutants (HTT-121, HTT-112 and HTT-101 and HTT-102) had higher CMTS than the heat-sensitive mutants (HTT-1 and HTT-105). Super Basmati had the lowest CMTS followed by HTT-1.

### Heat sensitive mutants have elevated level of ROS

ROS is one of the major byproducts of temperature stress and induces oxidative stress to plant. The higher MDA level in heat sensitive mutants under HTS give rise to a hypothesis that it might be due to increased ROS level. We thus measured *H_2_O_2_* from selected heat tolerant and sensitive mutants along with cv. Super basmati and sensitive check IR-64 at seedling stage. *H_2_O_2_ assay* indicated that although ROS was higher under HTS in all the tolerant and sensitive mutants, however, *H_2_O_2_* was accumulated more significantly in the sensitive mutants and moderately heat tolerant genotypes (super basmati and IR—64) as compared to tolerant mutants (Figure 7A). This indicated a more oxidative damage to these sensitive genotypes (HTT-1, HTT-105, super basmati and IR-64) as compared to tolerant mutants, thus leading to susceptibility towards heat stress. These results further suggested that higher *H_2_O_2_* level in the heat sensitive mutants could be due to the lower activities of antioxidant enzymes particularly CAT.

**Figure 7.**
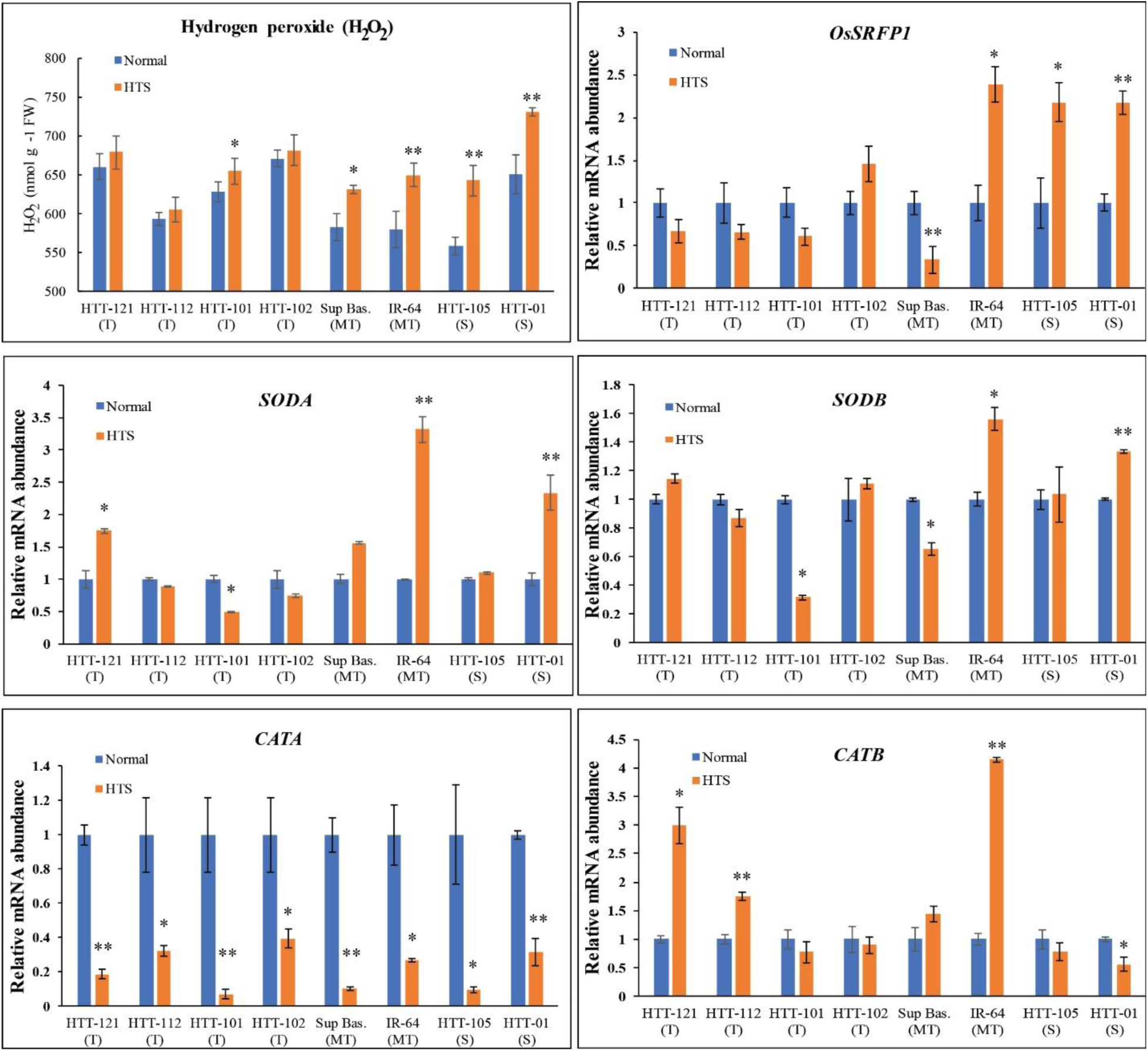
Hydrogen peroxide accumulation and relative expression analysis of stress responsive genes in contrasting heat tolerant mutants. (A) Quantification of H_2_O_2_ from contrasting heat tolerant mutants, Super basmati and IR-64. (B-F) Relative mRNA abundance of *OsSRFP1, SODA, SODB, CATA and CATB* genes in contrasting heat tolerant mutants, Super basmati and IR-64. Values indicate means of three biological replicates ± SD. Significance of data is tested by student’s *t* test. \**P*<0.05; \*\**P*<0.01.

### Relative expression of antioxidant genes

To further understand the underlying mechanism of heat tolerance in the tolerant mutants, we tested the expression level of few stress responsive genes particularly those involved in ROS producing and scavenging. *OsSRFP1* is known to negatively regulate abiotic stresses particularly cold, salt and oxidative stress via enhancing ROS level in rice (Fang *et al*. 2015; Fang *et al*. 2016). We observed a significant increase in the expression of *OsSRFP1* under HTS in the sensitive mutants and moderately sensitive cv. IR-64 (Figure 7B). This was consistent with the significantly increased ROS level in these mutants and IR-64 (Figure 7A) which points a possible positive correlation between them. This result indicated that *OsSRFP1* may negatively regulate heat stress tolerance in rice. We then tested the expression level of antioxidant related genes. The expression level of *SODA* was overall increased in both the tolerant and sensitive mutants with the most significant increase in IR-64 followed by HTT-1 (Figure 7C). *SODB* also showed more or less similar trend with the highest expression in IR-64 followed by HTT-1 under heat stress condition (Figure 7D). This could be due to the negative feedback mechanism of ROS. Since the activity of CAT was significantly increased in heat tolerant mutants and decreased in moderately heat tolerant varieties and sensitive mutants, we tested the expression level of *CATA* and *CATB* genes. *CATA* showed a significant decrease in the expression level under HTS in all the tested mutants along with cv. Super basmati and IR-64 (Figure 7E). However, *CATB* had significantly increased expression in the two most heat tolerant mutants (HTT-121 and HTT-112) but decreased expression in sensitive mutant, HTT-1 (Figure 7E). This was consistent with the CAT activity data which suggested that increased expression of *CATB* was involved in the increased activities of CAT enzyme under HTS in the heat tolerant mutants.

## Discussion

Paddy yield is a complex quantitative trait that is influenced by the genetic background of genotypes and environmental factors (Arshad *et al.* 2017; Wu *et al.* 2012). Mutants have been used to characterize genes for various important traits and to unravel physiological and molecular mechanisms of stress tolerance (Zhao *et al.* 2017). However, there have been no comprehensive evaluations of heat tolerance in rice mutants. Mutants have certain advantages over natural populations in underpinning genetic and physiological mechanisms for various phenotypic and physiological traits (Kurata *et al.* 2005). Recently, an EMS-mutagenized mutant of tomato, named *Slagl6*, was characterized for heat tolerance using genome editing, which improved our understanding of the mechanism of heat tolerance in tomato (Klap *et al.* 2017). In the present study, we developed a rice mutant population derived from cv. Super Basmati to identify useful heat-tolerant mutants to serve as an important resource for breeding and genetic studies. Mutants were evaluated under normal and HTS conditions to identify variation in important agronomic, physiological and biochemical traits at the seedling and reproductive stage. Earlier studies on rice were mostly conducted in growth chambers (at the seedling stage) or under artificial temperature stress (at the reproductive stage) to evaluate heat tolerance (Liu *et al.* 2018; Poli *et al.* 2013). We evaluated rice at both stages under realistic field conditions of HTS at the reproductive stage followed by confirmation under controlled conditions.

Rice growth is divided into three main developmental stages—vegetative, reproductive and ripening (http://ricepedia.org/rice-as-a-plant/growth-phases). Vegetative growth is mainly comprised of seedling development and active tillering followed by booting. The reproductive phase includes panicle development, anthesis and pollination. Rice crops are sensitive to high temperature at multiple growth stages, but booting and anthesis are the most critical stages that result in heavy yield losses due to sterility (Chaturvedi *et al.* 2017; Shah *et al.* 2011; Zafar *et al.* 2018). During the initial field evaluation in 2014, high temperatures at both the vegetative and reproductive stages were recorded in Multan (designated HTS treatment), relative to those in Faisalabad (normal condition) (Figure S1). The last two weeks of August were critical for anthesis and pollination, and there were frequent high-temperature episodes in the last two weeks of September (Figure S1). Differences of 2.7–4.7°C between the control and HTS treatments were observed at the start of anthesis, which were significant for evaluating rice for heat tolerance (Maruyama *et al.* 2013; Ps *et al.* 2017). An increase of 1°C above the threshold temperature may reduce grain yields in cereals by 4.1–10% (Wang *et al.* 2012). The ANOVA showed significant variation in the mutants under normal and HTS conditions for most of the evaluated traits including grain yield (Table 1). The effects of environment and G × E were also significant for most of the studied traits, except TGW and carotenoids (Table 1). Similarly, all mutants were evaluated for their response to various growth-related morpho-physiological and biochemical traits at the seedling stage (data for selected heat-tolerant and heat-sensitive mutants are in Figures 4 and 5).

Principal component analysis revealed how the different morpho-physiological and biochemical traits contribute to the variation in heat tolerance. In the biplots, traits on the opposite sides of PC1 and PC2 have a negative association (Font i Forcada *et al.* 2014). In the HTS environment, PTP, MDA, POD and esterase lie on the negative coordinate of PC1 and PC2 and have a negative association with other traits on the positive coordinate, including TGW and PY (Figure 1B), which was further confirmed by correlation analysis (Figure 2). Under HTS conditions, SOD, CAT, APX and RWC fell very close to PY (Figure 1B), illustrating an association of these seedling traits with yield, which was further confirmed by correlation analysis (Figure 2). In the biplot, HTT-121 was the furthest mutant from the origin under HTS and even normal conditions. HTT-121 was ranked the most heat-tolerant mutant, having the lowest HSI-GY (–27.92) and high grain yield, panicle fertility, thousand-grain weight and antioxidant enzyme levels under both normal and heat-stress conditions. Furthermore, LFW and LDW in the normal environment and SFW, SDW, RWC and SOD in the HTS environment had strong associations with TGW and yield, which support our hypothesis that seedling traits can be used as a selection parameter for heat tolerance. Previously, SOD and CAT were identified as useful indirect selection criteria for drought tolerance in wheat based on their strong positive correlations with grain yield (Afzal *et al.* 2017; Tabarzad *et al.* 2017).

The heat susceptibility index for grain yield (HSI-GY) has been frequently used as a reliable tool to characterize or screen crop germplasm for heat-tolerance ability (Aziz *et al.* 2018). Based on the 2014 grain yield performance in the field, we categorized the mutants and sensitive check IR-64 from tolerant to sensitive (Figure 3). To validate the agronomic performance of these mutants and the reproducibility of data, we re-evaluated selected tolerant and sensitive mutants along with IR-64 for grain yield in temperature-controlled field conditions. We used the four best-performing tolerant mutants and two least-performing sensitive mutants along with parent cv. Super Basmati and IR-64 for the second evaluation in 2016. We observed a similar trend in grain yield and panicle fertility in both years, which verified that the data from 2014 was reproducible and consistent (Figure 4A and 4B). In both years, HTS decreased grain yield in the sensitive mutants and Super Basmati and IR-64, but increased yield in the selected tolerant mutants (HTT-121, HTT-112 and HTT-101 and HTT-102). During reproductive growth, panicle fertility is the most sensitive trait affected by HTS in rice which directly affects final grain yield (Chaturvedi *et al.* 2017; Jagadish *et al.* 2007). In the present study, HTS significantly reduced panicle fertility (PF) in heat-sensitive mutants but had no significant effect on tolerant mutants (Figure 4B and 6), which indicates the activity of tolerance machinery in these mutants. Thus, PF could be used as a good selection criterion for heat tolerance in crops, particularly rice (Jagadish *et al.* 2007).

HTS significantly reduced LFW in sensitive mutants (HTT-1 and HTT-105) and Super Basmati, with no significant effect on heat-tolerant mutants (Figure 4C). Rather, HTT-112 and HT-101 showed a non-significant increase in LFW under HTS, which indicates its usefulness as an indirect selection criterion for heat tolerance. Leaf RWC has been used to evaluate plant water status under drought or heat stress (Saura-Mas and Lloret 2007). Heat stress decreased RWC in wheat and thus had a negative effect on plant homeostasis (Hameed *et al.* 2012). Here, HTS significantly reduced RWC in heat-sensitive mutants, relative to heat-tolerant mutants. Based on our findings, we suggest that higher RWC could be a useful indirect selection criterion for heat tolerance at the seedling stage.

Among the various responses for temperature stress, the antioxidant defense system is a quick response system that plays an important role in protecting plants from ROS damage (Wahid *et al.* 2007). Genotypic variation exists among germplasm for their potential to respond for ROS by activating antioxidant enzymes (Hussain *et al*. 2019). Genotypes that maintain higher antioxidant levels to detoxify ROS usually have smaller yield reductions under HTS (Mohammed and Tarpley 2009). In our study, SOD and CAT levels increased in heat-tolerant mutants under HTS, while the reverse was true for heat-sensitive mutants (Figure 5A and 5B). The increase in CAT was more significant than SOD, due to its involvement in heat tolerance by scavenging ROS. MDA is an indicator of ROS-mediated oxidative damage to plant cells (Cao and Zhao 2008). Consistent with the results for antioxidant enzyme levels, MDA levels increased more in heat-sensitive mutants under HTS than heat-tolerant mutants (Figure 5C), which supported previous findings (Hameed *et al.* 2012). ROS is an important trigger of cell death and leads to membrane lipid peroxidation (Hussain *et al*. 2019). To see if increased MDA level under HTS in the sensitive mutants is accompanied by higher ROS level, we measured *H_2_O_2_* from selected heat tolerant and sensitive mutants along with cv. Super basmati and sensitive check IR-64 at seedling stage. A significantly higher level of *H_2_O_2_* was observed in the heat sensitive mutants and moderately heat tolerant cv. Super basmati and IR-64 (Figure 7A) which was consistent with MDA data (Figure 5C). This leads to a suggestion that higher MDA level in the sensitive mutants was due to increased ROS accumulation.

To understand the molecular mechanism of increased ROS level and lower antioxidant activities in the heat sensitive mutants, we tested relative expression level of some stress responsive genes namely *SODA*, *SODB*, *CATA*, *CATB* and *OsSRFP1* (Fang *et al*. 2015; Fang *et al*. 2016; Zhao *et al*. 2018a; Zhao *et al*. 2018b; Das *et al*. 2019). *OsSRFP1* has been reported to be negatively involved in salt and cold tolerance in rice via positively regulating *H_2_O_2_* level (Fang *et al*. 2015; Fang *et al*. 2016). However, its role in heat tolerance has not yet been reported. In our study, we observed a significant upregulation in the expression of *OsSRFP1* in the heat sensitive mutants and sensitive check IR-64 which was consisted with increased *H_2_O_2_* level in these mutants (Figure 7B). This suggested its role as a negative regulator of heat stress tolerance in rice and seems that its role as a negative regulator of abiotic stresses is conserved. Rice *SOD* and *CAT* are key stress responsive genes which regulate the level of SOD and CAT enzymes in rice (Zhao *et al*. 2018a; Zhao *et al*. 2018b). Higher expression of these genes is linked with improved heat tolerance in rice (Zhao *et al*. 2018a; Zhao *et al*. 2018b; Das *et al*. 2019). In our study, we observed significant upregulation in the expression of *SODA* and *SODB* genes mainly in the heat sensitive mutants (Figure 7C,D). Since ROS is an important regulator of the expression of several genes (Mittler, 2017). We believe that this increased expression of *SODA* and *SODB* genes is due to increased ROS level in these mutants probably via feedback mechanism. Similarly, CATA gene also indicated a significant upregulation in all the tested mutants and control (Figure 7E). This could be due to negative feedback regulation of ROS. Notably, the expression of *CATB* gene was upregulated in the heat tolerant mutants and downregulated in the heat sensitive mutants. This is consistent with the CAT activity under heat stress, suggesting a key role of *CATB* in heat tolerance in rice. *CATB* has also been reported for its potential role in heat tolerance in rice at reproductive stage (Zhao *et al*. 2018a). Here we show that it is also involved in heat tolerance at early growth stages.

Based on these results and previous reports (Jagadish *et al.* 2010; Scafaro *et al.* 2010; Zafar *et al.* 2018), it could be inferred that the high antioxidant levels under HTS resulted in ROS scavenging, which played a key role in heat tolerance. Thus, higher SOD and CAT activities and lower MDA levels could serve as important selection indices for heat tolerance at early growth stages in rice. Since high temperature results in water loss from plant tissues, it exerts a negative pressure on the cell membrane and causes loss of cell turgidity. The stability of cell membranes thus decides the level of injury to plant cells and organelles. Higher cell membrane stability is therefore an indicator of drought and heat tolerance (Rehman *et al.* 2016). Our findings agree with previous studies, as we observed lower CMTS in heat-sensitive mutants than heat-tolerant mutants under HTS (Figure 5D).

Based on our findings, mutant HTT-121 was the most heat-tolerant and performed well under HTS. In contrast, HTT-1 was the most heat-sensitive mutant with poor performance for seedling-based morpho-physiological and biochemical traits as well as yield-related agronomic traits. Importantly, various seedling-based morpho-physiological (LFW, RWC, CMTS and MDA) and biochemical (SOD, CAT and H_2_O_2_) traits had strong positive associations with higher grain yield and could be used as selection criteria for heat tolerance in rice at early growth stages. PTP had a negative correlation with yield-related traits. Thus, high tiller number is not a good selection trait when breeding for high yield. Although, the role of *OsSRFP1* as a negative regulator of salt and cold stress has been reported previously, but we highlight for the first time the role of *OsSRFP1* in regulating heat tolerance in rice. Further studies, using overexpression and knockdown approaches could further strengthen these findings. Furthermore, the heat-tolerant mutants HTT-121, HTT-112, HTT-101 and HTT-102 could serve as a potential resource for developing mapping populations in further studies, especially quantitative trait loci mapping and map-based cloning of candidate genes related to higher yield under elevated temperature.

## Acknowledgments

The work was financially supported by the International Atomic Energy Agency (contract no. 16589). SAZ won a scholarship from Government of Punjab, Pakistan, for financial support during studies. We thank Dr. Saeed Rauf, University College of Agriculture, University of Sargodha, and Dr. Awais Rasheed, International Maize and Wheat Improvement Centre (CIMMYT) c/o CAAS China, for their generous assistance in data analysis and manuscript proof-reading. We sincerely acknowledge Pakistan Meteorological Department for providing the data of temperature.

